# Microfluidic Isolation of Aptamers with Affinity towards Multiple Myeloma Monoclonal Immunoglobulins (M-Ig)

**DOI:** 10.1101/2020.09.22.308650

**Authors:** Timothy R. Olsen, Claudia Tapia-Alveal, Milan N. Stojanovic, Tilla S. Worgall, Qiao Lin

## Abstract

Multiple myeloma (MM) is a bone marrow cancer of resident plasma cells that affects 125,000 patients in the U.S. with ∼ 30,000 new cases per year. Its signature is the clonal over-proliferation of a single plasma cell that secretes a patient specific monoclonal immunoglobulin (M-Ig). Detecting this patient specific M-Ig could allow sensitive detection of minimal residual disease in multiple myeloma from patient serum. Aptamers, single-stranded oligonucleotides with affinity and specificity to a target molecule, have recently been introduced as affinity reagents able to detect MM M-Igs. Here we adapt these benchtop M-Ig systematic evolution of ligands through exponential enrichment (SELEX) techniques to our bead integrated microfluidic SELEX (BIMS) device to rapidly generate patient specific aptamers. Using MM patient serum, we isolate patient M-Ig specific aptamers rapidly (runtime < 12 hours) with high affinity (K_D_ < 20 nM) while consuming limited quantities of patient M-Ig (< 100 μg).

## 1. Introduction

Multiple myeloma (MM) is a cancer of plasma cells primarily found in the bone marrow (BM) that affects nearly 96,000 in the U.S. ^1^. The disease is preceded by monoclonal gammopathy of undetermined significance (MGUS) that is asymptomatic but requires monitoring^2-3^. In MM a single plasma cell proliferates into a large clonal population that secrete monoclonal immunoglobulins (M-Ig) with identical variable regions. M-Ig are patient and tumor-specific and correlate with tumor burden^4-6^. The goal of treating MM is to achieve complete response (CR), defined as the absence of the hallmark monoclonal immunoglobulin (M-Ig) protein by immunofixation and less than 5% of cells as plasma cells in bone marrow (BM)^7^. Of patients who achieve CR, those who are negative for minimal residual disease (MRD), i.e., the presence of small numbers of myeloma cells in a patient’s body ^8^, have better survival than those who are MRD positive ^9^. Detection and monitoring of MRD, and thus the M-Ig, is hence emerging as a cornerstone in selecting and guiding therapeutic strategies^10^.

The two leading MRD diagnostic technologies, multiparameter flow cytometry (MFC) and VDJ sequencing (NGS), require plasma cells from bone marrow (BM) aspirates^9, 11-15^. Although both methods are very sensitive they have shortcomings including the non-representative nature of aspirates due to suboptimal procedures, the lack of uniform BM infiltration; and the poor viability of plasma cells after aspiration. BM biopsies are moreover costly, painful and not applicable to evaluation of extramedullary disease^16-20^. All of the above prevent frequent repeats and increase the likelihood of false-negative results^14, 21^. Recent studies have also explored the detection of MM through peptides that detect plasma cell released exosomes but these studies have not been applied to the detection of MRD^22^.

We recently demonstrated a method for sensitively detecting the presence of a patient M-Ig in serum using aptamers as an affinity receptor targeting the antigen binding fragment (fab) of the M-Ig^23^. We isolated aptamers that specifically recognized the idiotype and allowed low concentrations of M-Ig (<4.5 μg/dl M-Ig in serum detected) to be detected by effectively rejecting the effects of unspecific background proteins. We also recently developed a microfluidic device (termed BIMS) that integrates the aptamer selection process (SELEX) into a single platform without relying on multiple chips, and offline procedures^24^. The device combines highly efficient microfluidic reactions with a bead-based protocol to drastically reduce the time and rounds needed to isolate aptamers. BIMS was applied to the isolation of aptamers targeting gangliosides and could identify aptamers within a day while using small amounts of target (<200 μg).

While our M-Ig aptamer isolation protocol could reliably isolate aptamers, it utilized benchtop processes without automation. Here we present a method to rapidly generate specific and high affinity M-Ig targeting aptamers by adapting our BIMS device demonstrated previously to our M-Ig SELEX protocol. The two approaches are combined and further enhanced by the application of next generation sequencing into the aptamer selection analysis procedure for a more comprehensive assessment of the SELEX process. We join the two approaches to offer a method that can isolate aptamers rapidly while using modest amounts of patient samples. These qualities would allow isolation of patient specific aptamers for individuals that have been diagnosed with MM. Ultimately, a high affinity aptamers (Kd < 100 nM) that can be used in an aptamer formatted enzyme linked immunoassay (ELISA) to detect the presence of the cancer patient’s M-Ig is rapidly isolated (within a day) while consuming only modest amounts of a patient’s serum sample (<100 μg).

## 2. Materials and Methods

### Materials

MgCl_2_, human polyclonal IgG and molecular biology grade water were purchased from Sigma-Aldrich (St. Louis, MO). Deoxyribonucleotide triphosphates (dNTPs) and GoTaq Flexi DNA polymerase were obtained from Promega Corp. (Madison, WI). Protein G ELISA plates, Dulbecco’s phosphate buffered saline (D-PBS), Melon Gel IgG Spin Purification kits, GeneJET Purification Kits, Zeba Desaling Columns and streptavidin coupled agarose beads (Pierce Streptavidin Agarose) were purchased from ThermoFisher (Waltham, MA). Streptavidin surface plasmon resonance chips were purchased from Nicoya Lifesciences (Kitchener, ON, Canada). MiSeq 300 v2 kits and PhiX Control were purchased from Illumina (San Diego, CA). Next generation sequencing Kappa Hyper Prep Kits were purchased from Roche (Basel, Switzerland). AMPure XP magnetic purification beads were purchased from Beckman Coulter (Pasadena, CA). Randomized oligonucleotide library (5’ – TGC CAG CAT CGT AAT AGC CTC – 40N – ACC AAG TGA ATG AGC GGT ACG– 3’) and primers (Forward Primer: 5’ – TGC CAG CAT CGT AAT AGC CTC −3’, and Reverse Primer: 5’ – CGT ACC GCT CAT TCA CTT GGT – 3’) were synthesized and purified by Integrated DNA Technologies (Coralville, IA). Multiple myeloma patient serum was acquired from Tilla Worgall. Protein G beads were purchased from Santa Curz Biotechnology (Dallas, TX). Barcodes with Illumina adapters were purchased from Bioo Scientific (Austin, TX).

## Methods

### Selection of Aptamers towards Patient M-Protein

Patient serum was purified with Melon Gel following the manufacturer’s protocol. The melon gel binds to all serum proteins except IgG allowing extraction of a patient’s IgG. The extracted IgG was suspended in selection buffer (PBS with 2 mM MgCl_2_) and measured by UV-Vis (Implen P300, Implen GmbH, Munich, Germany). Protein G beads were rinsed three times with selection buffer and incubated with the patient IgG for 30 minutes. The beads were then washed five times with selection buffer. Extracts of this process were collected and used to verify successful conjugation by polyacrylamide gel electrophoresis (PAGE). Similarly, counter selection beads were prepared by incubating polyclonal IgG with washed protein G beads for 30 minutes and then washing the beads five times with selection buffer.

With selection beads prepared, six rounds of the SELEX process were performed as described in our previous work with some modifications. First, 20 pmoles of library were used to initiate the SELEX process. In the final two rounds (five and six) the washing time was increased from 35 minutes (700 μL) to 45 minutes (900 μL) to increase the selection pressure on these later rounds of the process (Table 1). Finally, a fraction of the eluted product of each round was collected for analysis by next generation sequencing (NGS).

**Table 1:** Selection parameters for patient monoclonal protein SELEX.

| Round | Wash Flow Rate | Wash Time | Counter Selection |
| --- | --- | --- | --- |
| 1 | 20 μL/min | 35 min | None |
| 2 | 20 μL/min | 35 min | Protein G Beads |
| 3 | 20 μL/min | 35 min | Polyclonal Beads |
| 4 | 20 μL/min | 35 min | Polyclonal Beads |
| 5 | 20 μL/min | 45 min | Polyclonal Beads |
| 6 | 20 μL/min | 45 min | Polyclonal Beads |

### Surface Plasmon Resonance (SPR)

SPR was used to discern the binding properties of the M-Ig towards aptamer candidates using a BioNavis Naali (BioNavis, Finland) instrument. Biotinylated aptamers (500 nM) were injected over a streptavidin chip at a flow rate of 20 uL/min for 5 minutes. After allowing the baseline to settle, a range of concentrations of M-Ig were injected. Surfaces were regenerated with 1 M sodium acetate between injections. Specificity was confirmed by injecting the polyclonal IgG over the sensor surface. Data from all injections was analyzed by 1:1 model fitting with BIAevaluate (Biacore, Slovakia).

### High Throughput Sequencing

Aptamer candidates from each round were sequenced using a MiSeq platform (Illumina, San Diego, CA). First the collected products of each round were amplified off-chip by PCR and checked with a gel electropherogram. The amplified product was purified and suspended in molecular biology grade water with a GeneJet DNA clean up kit to remove primers and the amount of DNA was quantified in the resulting solution using an Implen Spectrophotometer (Implen GmbH, Munich, Germany). 250 ng of the amplified product was used with a Kapa Hyper Prep kit and Bioo Scientific barcodes to ligate Illumina adapters. A unique barcode (6 total) was ligated to the product of each round of the SELEX allowing sequences from each round to be discernible in the NGS read file. The ligated product was verified to be of the correct length and to be of high purity by gel electrophoresis and the amount of DNA was quantified with an Implen spectrophotometer.

The barcoded samples were sequenced in two sequencing reactions. The first sequencing process sequenced rounds one to four, and a second sequencing process was used to sequence rounds five and six. Both reactions were prepared with the following procedure. A MiSeq 300 v2 cartridge was thawed to room temperature. Meanwhile, the barcoded samples were combined and diluted to create a 4 nM and 5 μL combined sample pool equally represented by each SELEX round (rounds 1-4 or rounds 5 and 6). 5 μL of 0.2 N NaOH was added to the 4 nM combined pool, mixed and left to incubate for 5 minutes at room temperature. A 1 mL and 20 pM solution was then prepared by adding 990 μL of the HT1 buffer provided by Illumina. A 600 uL and 4 pM solution was then prepared by withdrawing 120 uL from this solution and adding 480 μL of HT1 buffer. PhiX control was similarly diluted in HT1 buffer to produce a 4 pM solution of PhiX. The 4 pM PhiX (120 μL) and combined pool were mixed together to form a 600 μL solution. This solution was injected into the Illumina MiSeq cartridge and sequenced with an Illumina MiSeq instrument. Aptasuite version 0.8.9 was used to evaluate the reads^25^. From this software the cluster, and sequence analysis modules were used to determine which sequences should be ordered for in vitro testing.

### Enzyme-Linked Immunoassay (ELISA)

Aptamers were screened for their ability to bind to the monoclonal using an aptameric ELISA. The ELISA format consisted of monoclonal protein bound to the wells through protein G which then had a solution phase biotinylated aptamer applied. An increase in the absorbency intensity was indicative of aptamers binding to the target monoclonal protein. The wells were first washed three times with wash buffer (PBS with 2 mM MgCl2 added). Then 2.5 pmoles of target IgG or polyclonal IgG diluted in SuperBlock blocking buffer with 0.05% tween added were applied to the wells. The protein was incubated in the wells for 30 minutes at room temperature on a rotator. These wells were then washed eight times (∼200 μL per well per wash) with wash buffer. A 25 pmol solution of biotinylated aptamers in 100 μL of wash buffer were incubated in the washed wells for 60 minutes on a rotator. The wells were washed ten times (∼200 μL per well per wash) with wash buffer following the aptamer incubation. Following the wash, streptavidin horseradish peroxidase was diluted according to the manufacturer’s instructions in wash buffer and bovine serum albumin (BSA) and then applied to each well and allowed to incubate for 30 minutes at room temperature with an opaque cover. The wells were then washed 12 times with wash buffer. Finally, freshly mixed tetramethylbenzidine and substrate (1:1) were added to each well and the absorbance was recorded by an absorbance plate reader (VersaMax, Molecular Devices, San Jose, CA). Measurements were made at 450 nm in ten minute intervals.

## 3. Results and Discussion

### Patient M-Ig Selection

We developed a protocol to isolate aptamers for M-Igs directly from MM patient serum. Prior to treatment, the M-Ig will be highly abundant (>90% of sera IgG) in the patient serum, and therefore can be used as a target for aptamer selection. We designed our approach around the concept that we would collect serum from an MM patient, purify the M-Ig from the serum (using antibody specificity), and then inject that purified antibody into the BIMS device to generate M-Ig specific aptamers (**Figure 1**).

**Figure 1:**
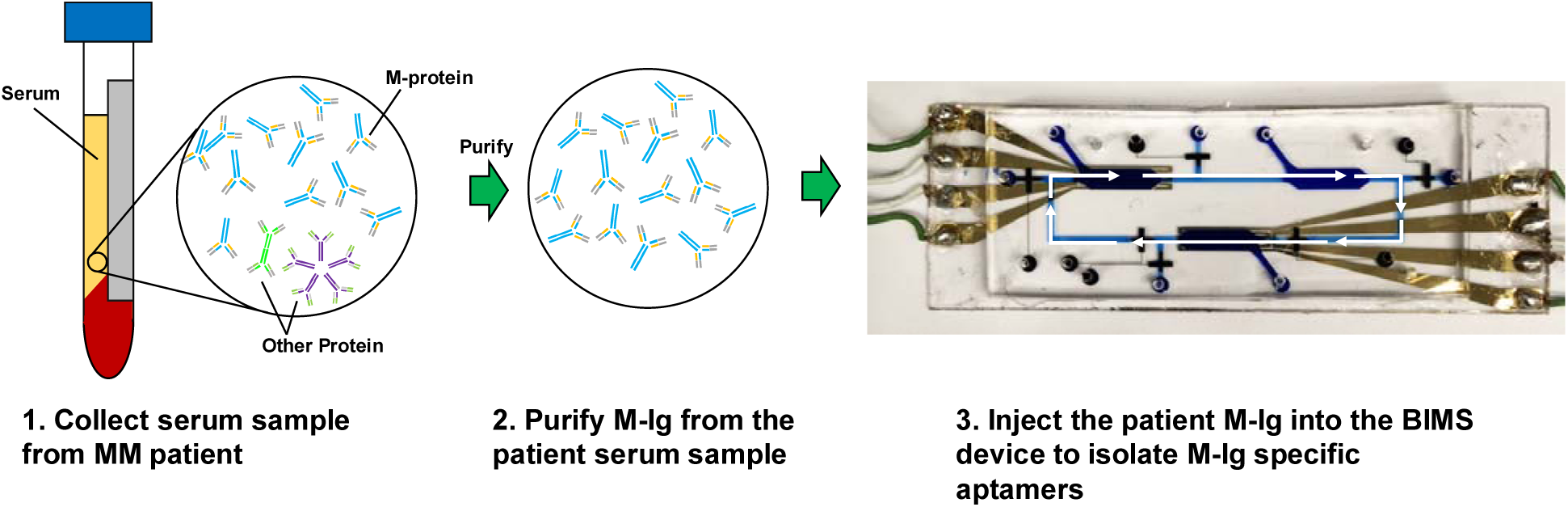
Aptamer selection process for a multiple myeloma patient. First the patient serum sample is conditioned to remove non-IgG proteins. The remaining IgG proteins are then bound to beads and used as the target in the microfluidic SELEX process.

A myeloma patient’s serum sample was first prepared for the SELEX. Patient serum was purified with Melon Gel. The melon gel binds to all serum proteins except IgG allowing extraction of a patient’s IgG. Because of the nature of IgG myeloma, the extracted IgG from the patient’s serum will be dominated by the monoclonal IgG. The purified protein was then bound to Protein G beads and used as a target in the BIMS device, while polyclonal IgG were also prepared and used as a counter target.

With selection beads prepared, we performed six rounds of the SELEX process. This approach differed from our previous work in several ways. First, a more concentrated library was used to initiate the SELEX process. A greater amount of starting library amount was used to increase the potential to isolate aptamers with very high specificity. Second, in the first round of SELEX counter selection beads were not used. This was to reduce the potential for losing high affinity sequences that are present at low abundance, as would be expected from library members. The second round used unreacted protein G beads as counter targets. Rounds three to six used polyclonal IgG beads as a counter target. Further, in the final two rounds (five and six) the we used longer washing durations to increase the selection pressure on these later rounds of the process. Finally, a fraction of the eluted product of each round was collected for analysis by next generation sequencing (NGS).

The success of the overall chip performance was first assessed using eluates collected during the selection process. These eluates were amplified off-chip by PCR and imaged by gel electrophoresis. As can be seen in **Figure 2**, for every SELEX round the amount of DNA collected becomes less as the washing progresses. However, after the thermal conditions of the selection chamber are changed, a large amount of DNA that was strongly bound and eluted under the different environmental conditions could be collected. Each round of SELEX showed the monotonous decrease in the amount of DNA during washing. The first wash of the first round of selection appears to have a lower intensity but is actually artifact of having an excess of DNA in the PCR reaction. All samples were subjected to the same amount of PCR cycles (19 cycles). Later rounds of SELEX had more oligonucleotides in the elution compared to even the first wash indicating more of the pool was binding to the target beads than being washed away during the later rounds of SELEX.

**Figure 2.**
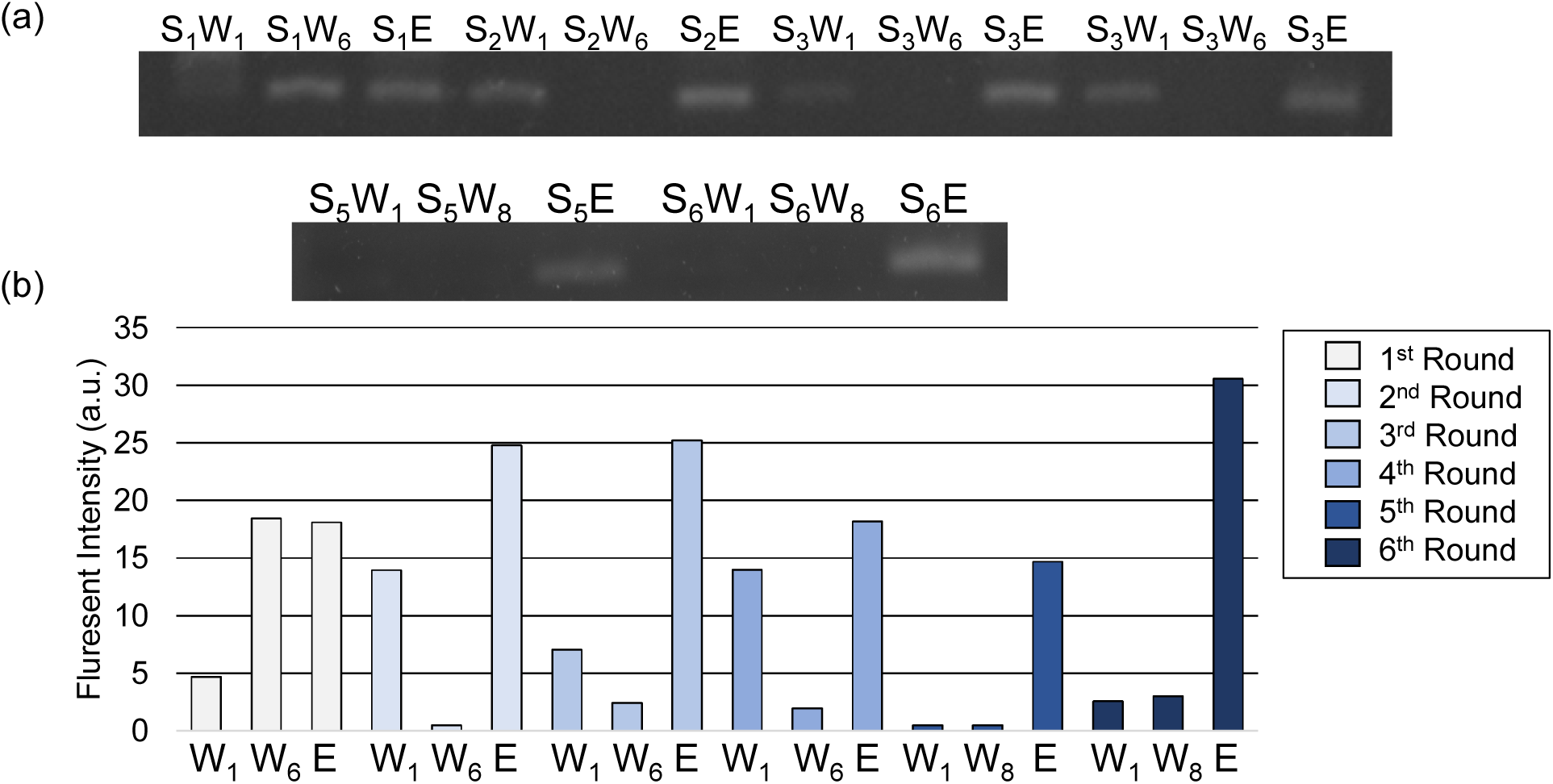
Collected eluates from the six round selection process, amplified by PCR and imaged with gel electrophoresis. (a) Gel electropherogram of amplified eluates and (b) bar graph representing electropherogram band intensity. Bar color indicates selection round. W_1_ is the first wash of a selection round. W_6_ is the final wash for section rounds 1-4. The final wash for rounds 5 and 6 is W_8_. E is the eluted product.

### Sequence Analysis

We collected eluate from the BIMS device and sequenced with next generation techniques and analyzed with the AptaSuite toolkit ^25^. The toolkit fractioned the raw sequencing files into the six rounds of SELEX and accepted 3,818,436 reads as having correct forward primers and reverse primers. Of these reads, approximately all (>95%) had a randomized region size matching the starting library (40 nucleotides). The identified barcodes for each round of SELEX were approximately evenly distributed (first round: 19.1%, second round: 18.38%, third round: 17.31%, fourth round: 13.31%, fifth round: 16.12%, and sixth round: 15.57%). The amount of singletons, defined as sequences that appear with a count of one, decreased on every round of SELEX (**Figure 3**). Further, the amount of enriched species, defined as sequences that also appear in other rounds of SELEX increased as expected during every round of SELEX. Conversely, the unique fraction (sequences that only appear in that SELEX round) decreases as the SELEX process progressed. It appears the singletons and enriched sequences slightly increased on the sixth round. This could indicate the process began to saturate after six rounds.

**Figure 3.**
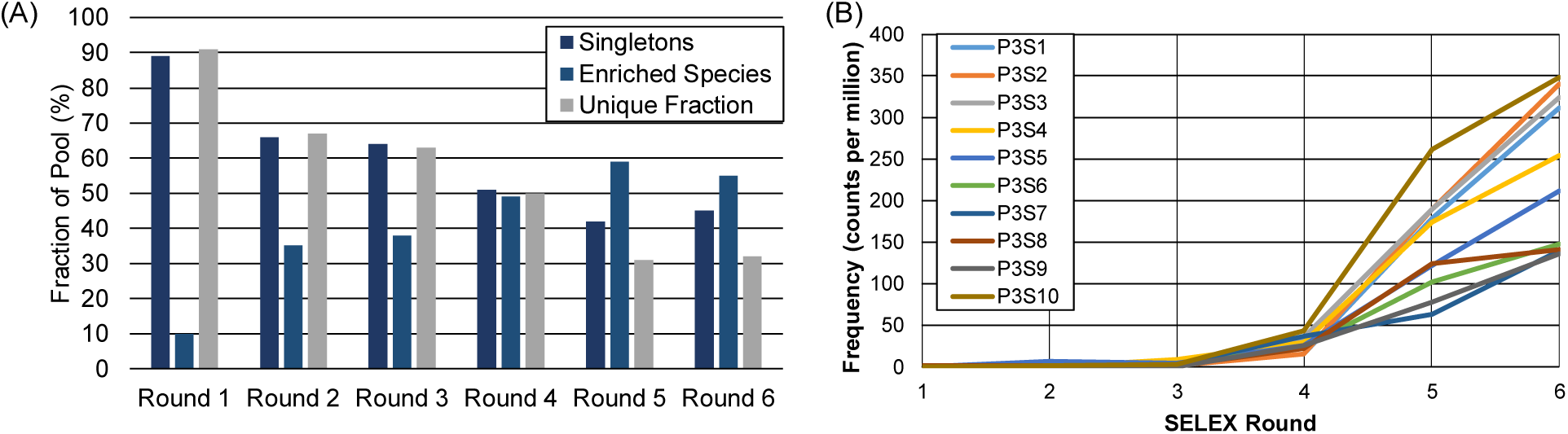
(A) Singletons, enriched species and unique fraction of every SELEX round. Singletons are sequences that appear with a frequency of one. Enriched species are sequences that are found in other rounds. The unique fraction are the sequences that do not appear in any other round. (B) Abundance of the top ten most frequent sequences from the sixth round of SELEX. Significant enrichment occurs from the fourth round to the fifth round.

A cluster program was run to group aptamer sequences by their similarities and the clusters of similar sequences. Clusters consisted of sequences that had randomized regions with only a single nucleotide allowed to mismatch. Clusters that appear at very high frequencies in the sixth round of SELEX were further evaluated for their progression through the other rounds of SELEX. A sequence of low abundancy but that is rapidly increasing in frequency in the later rounds of SELEX could suggest a potential high affinity sequence becoming enriched. The clusters were checked against the most frequent non-clustered sequences to ensure highly frequent sequences were not missed by the clustering algorithm. Sequences that contained a single nucleotide between primers were considered artifacts from the PCR processes and purification processes and discarded. The progression of the ten most abundant sequences from the sixth round are shown **Figure 3**. Their sequences and abundance values are tabulated in Table 2.

**Table 2:**
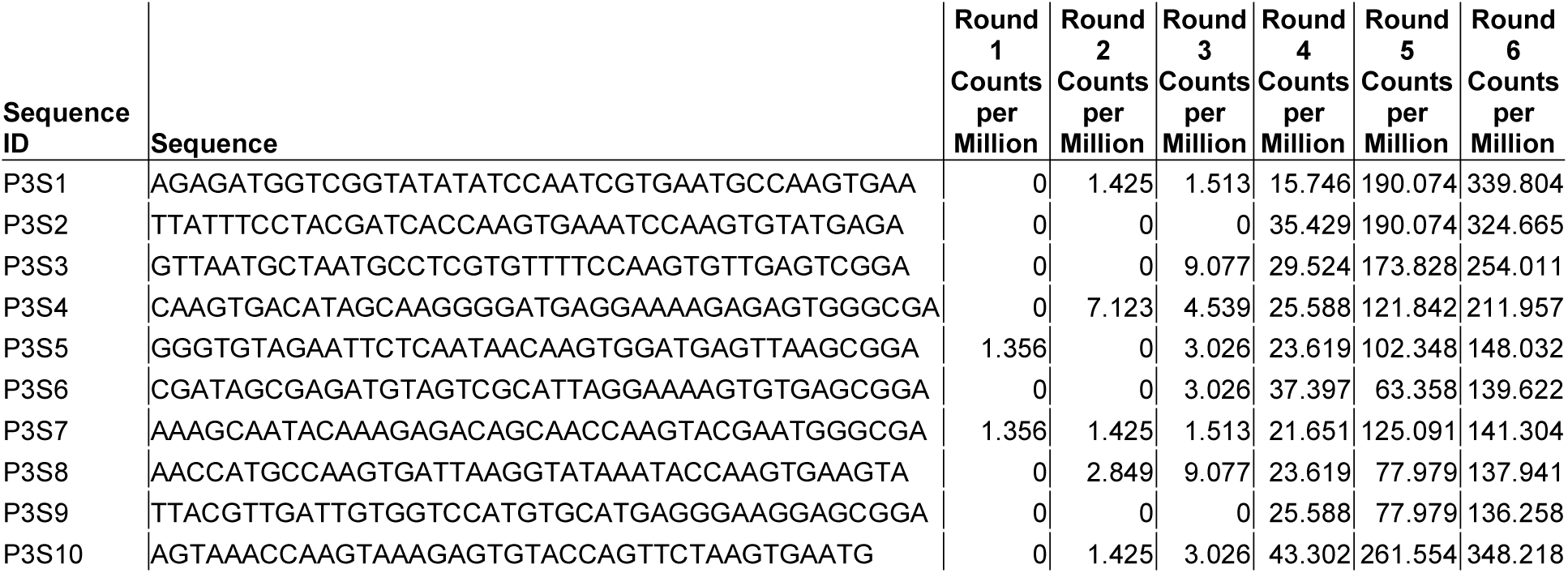
Next generation sequencing results for each round of SELEX for the ten most frequent sequences found in the 6^th^ round of SELEX.

All of the most frequent sequences increased from the fifth round to the sixth round. The most significant enrichment of the sequences occurred after the fourth round indicating that SELEX cycles beyond four could enhance the probability of identifying a high affinity aptamer. The most frequent sequences of the fourth and sixth SELEX rounds were synthesized for further in vitro testing. These sequences are tableted in the supplementary information.

### Affinity and Specificity of Aptamers

We took these ten most prevalent sequences and screened them with an ELISA **(Figure 4**). All sequences except P3S2, P3S7, and P3S8 showed significantly greater binding towards the M-Ig than the polyclonal. Of these, it was observed P3S10 and P3S5 had the greatest difference in binding towards the M-Ig than the polyclonal.

**Figure 4:**
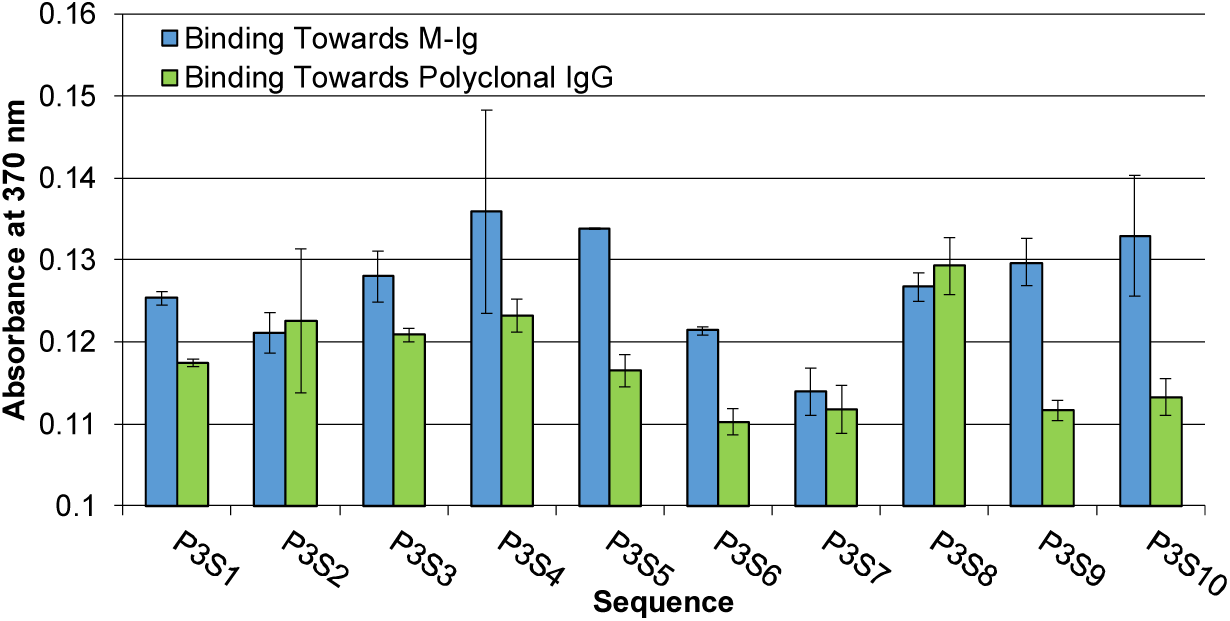
Aptameric ELISA results of a patient monoclonal IgG bound to the well surface through protein G and solution phase aptamer. Absorbance was measured at 370 nm in 10 minute intervals. Measurement shown is after 75 minutes. Error bars represented standard deviations from triplicate measurements.

These results can be compared to their enrichment found through NGS where sequence P3S10 was the most prevalent sequence for the fourth, fifth, and sixth SELEX round. While it was the most prevalent for the sequences tested, there were other sequences at the fourth round which were more frequent than P3S10 but these sequences became less frequent in the fifth and sixth round.

We further analyzed the aptamer with greatest binding from the ELISA (P3S10) with by SPR (**Figure 5**). A biotinylated version of the aptamer was bound to one of the sensing surfaces while a scrambled control was bound to the other. SPR measurements were taken from the difference of these two channels such that only the specific binding of the aptamer to the injected molecule was considered. Patient serum purified for IgG by Melon Gel was injected into the SPR instrument in various concentrations. The aptamer and scrambled sequence were then regenerated and human polyclonal IgG was then injected at the same concentrations and compared to the patient injections. These experiments were performed in triplicate for both the M-Ig and polyclonal IgG.

**Figure 5:**
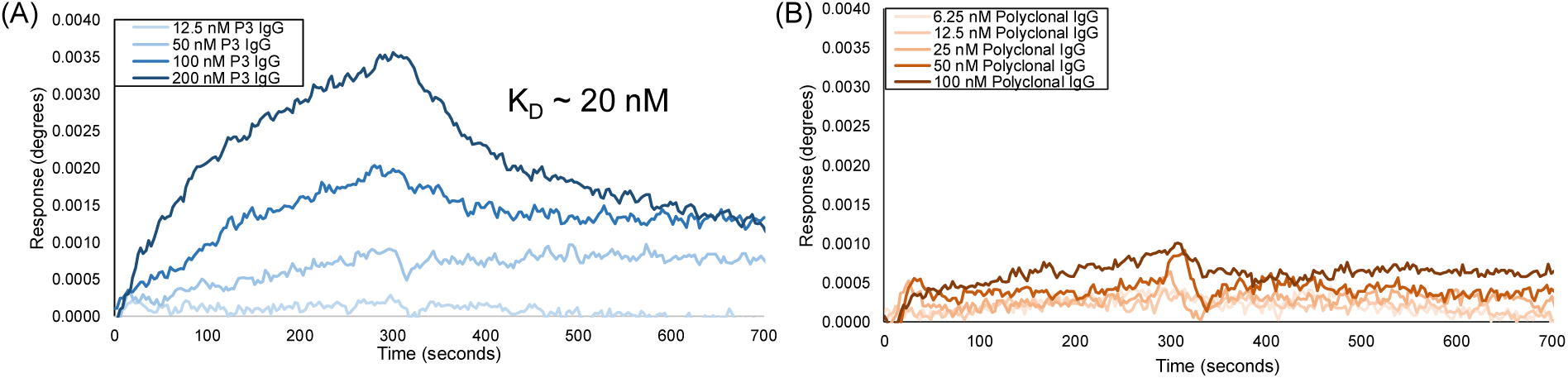
SPR response curves for aptamer P3S10. A binding dissociation constant (K_D_) was found to be ∼20 nM

From the SPR responses it can be observed that significant binding was found to the patient IgG protein while negligible binding was found when the polyclonal human IgG was injected. The SPR plots exhibit two-phase events at high concentrations, where there are two regions of exponential decay. Each IgG molecule bound to the SPR surface has two potential Fab region binding sites that the aptamer can bind, thus we suspect aptamers are rebinding after dissociation which artificially lowers the dissociation rate constant. We applied a 1:1 fitting model to the M-Ig binding curves to the lower concentration injections which do not have two-phase dissociation to obtain an effective binding dissociation constant (K_D_) of ∼20 nM. This binding dissociation constant describes the avidity of the aptamers towards the multiple binding sites present on the target patient protein and includes the aptamer rebinding artificially lowering the k_off_. The concentrations of protein used in these SPR plots are orders of magnitude lower than the limit of detection for conventional MM diagnostics, though it must be noted the SPR proteins were diluted in buffer.

## 4. Conclusion

We applied our BIMS approach to the isolation of a patient derived monoclonal protein implicated in cancer. We directly took patient serum and isolated the cancer associated protein which was then put onto selection beads and directly injected into the microfluidic device. High throughput sequencing was used to analyze the resulting aptamer pools and to select sequences to be further analyzed. These sequences were analyzed by an aptameric ELISA and sequences P3S10 was found to have higher binding to the patient M-Ig than to the polyclonal control. P3S10 was further analyzed by SPR and found to have a specific response to the patient IgG and negligible response to a polyclonal IgG pool. The rapid isolation time of these aptamers (<12 hours) could allow the development of personalized probes for MRD diagnostics for a cohort of patients. These aptamers would be most useful in detection assays, especially MRD detection assays, where patient serum is combined with aptamers and a readout. Thus, we plan on characterizing these aptamers for their affinity and limit of detection in physiological solutions in future work.

## 5. Acknowledgements

This study was supported in part by the National Institutes of Health grants R21DK109690, R21CA199849, R33CA196470, TL1 TR001875, and UL1TR001873.

## References

1. Howlader, N.; Noone, A.; Krapcho, M.; Garshell, J.; Neyman, N.; Altekruse, S.; Kosary, C.; Yu, M.; Ruhl, J.; Tatalovich, Z., SEER Cancer Statistics Review, 1975–2012, National Cancer Institute; 2015. 2016.

2. Weiss, B. M.; Abadie, J.; Verma, P.; Howard, R. S.; Kuehl, W. M., A monoclonal gammopathy precedes multiple myeloma in most patients. Blood 2009, 113 (22), 5418–5422.

3. Landgren, O.; Kyle, R. A.; Pfeiffer, R. M.; Katzmann, J. A.; Caporaso, N. E.; Hayes, R. B.; Dispenzieri, A.; Kumar, S.; Clark, R. J.; Baris, D., Monoclonal gammopathy of undetermined significance (MGUS) consistently precedes multiple myeloma: a prospective study. Blood 2009, 113 (22), 5412–5417.

4. Kyle, R., Diagnostic criteria of multiple myeloma. Hematology/oncology clinics of North America 1992, 6 (2), 347–358.

5. Kyle, R.; Rajkumar, S., Criteria for diagnosis, staging, risk stratification and response assessment of multiple myeloma. Leukemia 2009, 23 (1), 3.

6. Kyle, R. A., The monoclonal gammopathies. Clin. Chem. 1994, 40 (11), 2154–2161.

7. van de Velde, H. J.; Liu, X.; Chen, G.; Cakana, A.; Deraedt, W.; Bayssas, M., Complete response correlates with long-term survival and progression-free survival in high-dose therapy in multiple myeloma. Haematologica 2007, 92 (10), 1399–1406.

8. Martinez-Lopez, J.; Lahuerta, J. J.; Pepin, F.; González, M.; Barrio, S.; Ayala, R.; Puig, N.; Montalban, M. A.; Paiva, B.; Weng, L., Prognostic value of deep sequencing method for minimal residual disease detection in multiple myeloma. Blood 2014, 123 (20), 3073–3079.

9. Rawstron, A. C.; Child, J. A.; de Tute, R. M.; Davies, F. E.; Gregory, W. M.; Bell, S. E.; Szubert, A. J.; Navarro-Coy, N.; Drayson, M. T.; Feyler, S., Minimal residual disease assessed by multiparameter flow cytometry in multiple myeloma: impact on outcome in the Medical Research Council Myeloma IX Study. Journal of Clinical Oncology 2013, 31 (20), 2540–2547.

10. Romano, A.; Palumbo, G. A.; Parrinello, N. L.; Conticello, C.; Martello, M.; Terragna, C., The minimal residual disease within the bone marrow of Multiple Myeloma: caveats, clinical significance and future perspectives. Frontiers in oncology 2019, 9, 699.

11. Korthals, M.; Sehnke, N.; Kronenwett, R.; Bruns, I.; Mau, J.; Zohren, F.; Haas, R.; Kobbe, G.; Fenk, R., The level of minimal residual disease in the bone marrow of patients with multiple myeloma before high-dose therapy and autologous blood stem cell transplantation is an independent predictive parameter. Biology of Blood and Marrow Transplantation 2012, 18 (3), 423–431. e3.

12. Tatsas, A. D.; Jagasia, M. H.; Chen, H.; McCurley, T. L., Monitoring Residual Myeloma High-Resolution Serum/Urine Electrophoresis or Marrow Biopsy With Immunohistochemical Analysis? Am. J. Clin. Pathol. 2010, 134 (1), 139–144.

13. Vij, R.; Mazumder, A.; Klinger, M.; O’Dea, D.; Paasch, J.; Martin, T.; Weng, L.; Park, J.; Fiala, M.; Faham, M.; Wolf, J., Deep Sequencing Reveals Myeloma Cells in Peripheral Blood in Majority of Multiple Myeloma Patients. Clinical Lymphoma Myeloma and Leukemia 2014, 14 (2), 131–139.e1.

14. Kumar, S.; Paiva, B.; Anderson, K. C.; Durie, B.; Landgren, O.; Moreau, P.; Munshi, N.; Lonial, S.; Bladé, J.; Mateos, M.-V., International Myeloma Working Group consensus criteria for response and minimal residual disease assessment in multiple myeloma. The Lancet Oncology 2016, 17 (8), e328–e346.

15. Martinez-Lopez, J.; Lahuerta, J. J.; Pepin, F.; González, M.; Barrio, S.; Ayala, R.; Puig, N.; Montalban, M. A.; Paiva, B.; Weng, L., Prognostic value of deep sequencing method for minimal residual disease detection in multiple myeloma. Blood 2014, blood-2014-01-550020.

16. Wei, A.; Westerman, D.; Feleppa, F.; Trivett, M.; Juneja, S., Bone marrow plasma cell microaggregates detected by immunohistology predict earlier relapse in patients with minimal disease after high-dose therapy for myeloma. Haematologica 2005, 90 (8), 1147–1149.

17. Lee, N.; Moon, S.; Kim, S.; Hwang, S.; Park, H.; Bang, S.-M.; Lee, J.; Yoon, S.-S.; Lee, D., Analysis of CD138+ Plasma Cell Percentage in Bone Marrow Section Using Image Analyzer: Discrepancies of Bone Marrow Plasma Cell Count between Aspiration and Biopsy Section in Multiple Myeloma. Clinical Lymphoma, Myeloma and Leukemia 2015, 15, e96.

18. Bergen, H. R.; Dasari, S.; Dispenzieri, A.; Mills, J. R.; Ramirez-Alvarado, M.; Tschumper, R. C.; Jelinek, D. F.; Barnidge, D. R.; Murray, D. L., Clonotypic light chain peptides identified for monitoring minimal residual disease in multiple myeloma without bone marrow aspiration. Clin. Chem. 2016, 62 (1), 243–251.

19. Udd, K. A.; Spektor, T. M.; Berenson, J. R., Monitoring Multiple Myeloma. Clinical advances in hematology & oncology: H&O 2017, 15 (12), 951–961.

20. Brestoff, J. R.; Toland, A.; Afaneh, K.; Qavi, A. J.; Press, B.; Westervelt, P.; Kreisel, F.; Hassan, A., Bone Marrow Biopsy Needle Type Affects Core Biopsy Specimen Length and Quality and Aspirate Hemodilution. Am. J. Clin. Pathol. 2018.

21. Cogbill, C.; Spears, M.; Vantuinen, P.; Harrington, A.; Olteanu, H.; Kroft, S., Morphologic and cytogenetic variables affect the flow cytometric recovery of plasma cell myeloma cells in bone marrow aspirates. International journal of laboratory hematology 2015, 37 (6), 797–808.

22. Iaccino, E.; Mimmi, S.; Dattilo, V.; Marino, F.; Candeloro, P.; Di Loria, A.; Marimpietri, D.; Pisano, A.; Albano, F.; Vecchio, E., Monitoring multiple myeloma by idiotype-specific peptide binders of tumor-derived exosomes. Molecular cancer 2017, 16 (1), 159.

23. Caludia Tapia-Alveal, T. R. O., Tilla S Worgall, Personalized immunoglobulin aptamers for detection of multiple myeloma minimal residual disease in serum.

24. Timothy R. Olsen, K.-A. Y., Kechun Wen, Xin Meng, Tyler J. Stewart, Stacy Hall, Nicholas J. Steers, Ali G. Gharavi, Jan Novak, Milan N. Stojanovic,, and Qiao Lin, Automatized Isolation of Aptamers for Glycoconjugates.

25. Hoinka, J.; Backofen, R.; Przytycka, T. M., AptaSUITE: a full-featured bioinformatics framework for the comprehensive analysis of aptamers from HT-SELEX experiments. Molecular Therapy-Nucleic Acids 2018, 11, 515–517.

